# Genome assembly of the southern pine beetle (*Dendroctonus frontalis* Zimmerman) reveals the origins of gene content reduction in *Dendroctonus*

**DOI:** 10.1101/2024.05.08.592785

**Authors:** Megan Copeland, Shelby Landa, Adekola Owoyemi, Michelle M. Jonika, Jamie Alfieri, Terrence Sylvester, Zachary Hoover, Carl E. Hjelmen, J. Spencer Johnston, Bethany R. Kyre, Lynne K. Rieske, Heath Blackmon, Claudio Casola

## Abstract

*Dendroctonus frontalis*, also known as southern pine beetle (SPB), represents the most damaging forest pest in the southeastern United States. Strategies to predict, monitor and suppress SPB outbreaks have had limited success. Genomic data are critical to inform on pest biology and to identify molecular targets to develop improved management approaches. Here, we produced a chromosome-level genome assembly of SPB using long-read sequencing data. Synteny analyses confirmed the conservation of the core coleopteran Stevens elements and validated the *bona fide* SPB X chromosome. Transcriptomic data were used to obtain 39,588 transcripts corresponding to 13,354 putative protein-coding loci. Comparative analyses of gene content across 14 beetle and 3 other insects revealed several losses of conserved genes in the *Dendroctonus* clade and gene gains in SPB and *Dendroctonus* that were enriched for loci encoding membrane proteins and extracellular matrix proteins. While lineage-specific gene losses contributed to the gene content reduction observed in *Dendroctonus*, we also showed that widespread misannotation of transposable elements represents a major cause of the apparent gene expansion in several non-*Dendroctonus* species. Our findings uncovered distinctive features of the SPB gene complement and disentangled the role of biological and annotation-related factors contributing to gene content variation across beetles.

## Background

Bark beetles (subfamily Scolytinae) are common forest pests responsible for the annual loss of millions of conifers and other trees worldwide (1). The genus *Dendroctonus* (Latin for “tree killer”) includes several bark beetle species that can spawn large outbreaks and are capable of colonizing and overwhelming both weakened and healthy trees. Beside timber loss, the ecological repercussions of large-scale forest destruction caused by *Dendroctonus* include ecosystem degradation, hydrological instability, reduced carbon sequestration and loss of revenue associated with commercial and recreational use.

The southern pine beetle (SPB) *Dendroctonus frontalis* Zimmerman has historically been associated with the most severe bark beetle epidemics in southeastern U.S., leading to the loss of millions of hectares of managed and unmanaged conifer forests (2, 3). Pioneer SPBs use visual and chemical cues to locate host trees and, once landed, bore into the phloem of the trees, avoiding resin ducts with defensive chemicals (2, 4). Adult female beetles create S-shaped galleries where eggs are laid; larvae feed on the phloem within their immediate vicinity (5). Upon reaching maturity, the adult beetles find a new host tree and continue to propagate (5).

Under ideal conditions, a population could produce up to eight generations within one year, leading to a potential for rapid population increase in an impacted area (6). When female pioneer beetles find a new host tree, they release frontalin, an aggregation pheromone. The beetle’s pheromones and the host tree’s distress odors attract males and females of this species to the host tree. This attraction process is critical as the number of attacking beetles necessary to overcome the host tree’s defenses positively correlates with the host tree’s strength. Changing climate patterns, including warming temperatures and fluctuating precipitation patterns, together with a lack of effective, proactive management, have allowed an unprecedented northward range expansion, and SPB is now infesting pine forests of New England (7).

Traditional integrated pest management strategies for *Dendroctonus*, including population density surveys, outbreak prevention and treatment of affected areas are costly and pose logistic challenges over large areas (6, 8, 9). Given the geographical expansion of SPB and other *Dendroctonus* species coupled with their persistent outbreaks over historic geographic ranges, additional tools are needed to develop innovative strategies for the management of these bark beetles. Genome-wide analyses are critical to identify the genetic basis of species-specific adaptations, including the suite of phenotypes associated with the tree-killing habit of SPB and several other *Dendroctonus* species.

Changes in chromosome number and structure and in gene content are fundamental sources of genetic variation that underlie adaptive evolution. The genus *Dendroctonus* shows a rapidly evolving karyotype, with 2n=30 being the presumed ancestral chromosome number that is still retained in a few species, and 2n=12 being the smallest karyotype (5). Several species experienced lineage-specific fusions of autosomes and ancestral sex chromosomes, leading to the formation of neo-XY chromosomes (10). The extreme karyotypic variation in *Dendroctonus* is further supported by the presence of chromosome number changes between populations of the same species. For instance, the two *D. frontalis* morphotypes A and B exhibit the karyotypes 7AA+XY and 5AA+XY, respectively (11). The analysis of the first sequenced *Dendroctonus* genome, the MPB, revealed a gene content of ∼13,000, which is significantly lower than in most other beetles and insects (12, 13). This finding has been confirmed by the genome sequencing of the red turpentine beetle (RTB), *D. valens* (14). Intriguingly, two other sequenced bark beetle species, *Ips typographus* (European spruce bark beetle) (15), and *Hypothenemus hampei* (coffee berry borer) (16, 17) contained between19,000-23,000 genes. The more distantly related wood-boring species *Anoplophora glabripennis* (Asian Longhorned Beetle) and *Agrilus planipennis* (Emerald Ash Borer) shared a similarly high gene content (18).

However, the causes of the apparent gene content reduction in the genus *Dendroctonus* have not yet been investigated.

In this study, we generated a high-quality genome assembly and gene annotation resources for the southern pine beetle, *D. frontalis*, using a combination of long-read sequencing, HI-C scaffolding and high-throughput transcriptomic data. We found synteny conservation between SPB largest scaffolds and chromosomes from other species, including conservation of the putative SPB X chromosome with the MPB neoX chromosome. The comparative analysis of SPB and other beetle genomes revealed SPB- and *Dendroctonus*-specific gene gains and losses potentially associated with adaptations and an inflated gene count in several non-*Dendroctonus* beetles due to the erroneous annotation of transposable elements.

## Materials & Methods

### Biological material and nucleic acid extraction

SPB specimens were collected from infested loblolly pine trees in the Homochitto National Forest, MS (31° 21’ 16.152 “N, 90° 49’ 42.678” W), between September 29 and October 7, 2019 **(Table S1)**. Four females and three males were collected after eclosion and were stored frozen until DNA extraction was performed. Whole genomic high molecular weight (HMW) DNA was extracted from three female and two male pooled sample sets using the MagAttract High Molecular Weight kit (Qiagen, Valencia, CA) according to the manufacturer’s protocols with the addition of an extra wash step using the provided wash buffer. HMW Genomic DNA was collected from two additional pooled sample sets, one female and one male, using the Nanobind Tissue Big DNA Kit (Circulomics, Baltimore, USA) according to the manufacturer’s protocols.

RNA was obtained from 3 female, 4 male and 39 larvae at various developmental stages stored in either RNAlater or liquid nitrogen after collection and subsequently maintained at −80 until shipment for sequencing **(Table S1)**. Total RNA was isolated from whole beetles with TRI Reagent RT (Molecular Research Center Inc., Cincinnati, OH), RNA integrity was verified using gel electrophoresis and absorbance was measure at 260/280 and 230/280. cDNA was synthesized using SuperScript III Reverse Transcriptase (Invitrogen, Carlsbad, CA) according to manufacturer’s instructions at a concentration of 3000 ng/ml and used as a template for the RT-qPCR standard curve, constructed using a 5-fold dilution.

### Genome sequencing

Long-read sequences were obtained utilizing the SQK-LSK109 reaction kit, and libraries were prepared following the manufacturer’s protocol. One R9.4.1 flow cell was used for each specimen, and base-calling was performed with Guppy (v3.2.10) using default system settings. Sequencing yielded a total of 39 Gb of read data (∼198X coverage).

### Omni-C sequencing

Dovetail Genomics performed the preparation of Omni-C libraries and sequencing (Dovetail Genomics, CA). For each Dovetail Omni-C library, chromatin was fixed in place with formaldehyde in the nucleus. Fixed chromatin was digested with DNaseI and then extracted. Chromatin ends were repaired and ligated to a biotinylated bridge adapter, followed by proximity ligation of adapter-containing ends. After proximity ligation, crosslinks were reversed, and the DNA was purified. Purified DNA was treated to remove biotin that was not internal to ligated fragments. Sequencing libraries were generated using NEBNext Ultra enzymes and Illumina-compatible adapters. Biotin-containing fragments were isolated using streptavidin beads before PCR enrichment of each library. The library was sequenced on an Illumina HiSeqX platform to produce approximately 30x sequence coverage.

### Genome Size Estimation

Flow cytometric methods following (19) were used to determine the D. frontalis genome size. Neural tissue from individual frozen samples of D. frontalis was dissected and deposited into 1 mL of Galbraith buffer. All samples were co-prepared with a standard (lab stock of *Drosophila virilis*, genome size = 328 Mbp). Samples were gently ground with a Kontes “A” pestle approximately 15 times to release nuclei. After passing samples through 41-micrometer mesh filters, samples were stained with 25 µl of 1mg/µl propidium iodide and incubated in the dark. Samples were run on a Beckman Coulter CytoFlex flow cytometer with a 488 nm blue laser. Means of 2C nuclei fluorescence peaks were measured for both sample and standard using gating methods supplied within the instrument’s software before calculating the estimated genome size.

### Genome assembly

Initial assembly was built using all female sequencing reads (approximately 19 Gb of reads) in CANU version 1.9 (20). The estimated genome size was set to 200 Mb based on flow cytometry estimates. Nanopolish version 2.5.0 was used to improve the consensus sequence using standard settings. This process yielded a final assembly with 1436 sequences, a total length of 209 Mb, and an N50 of 1.61 Mb.

We assembled the female SPB genome using all female reads (approximately 19 Gb of reads) in flye version 2.8.2-b1689 with default settings (21). We then used Blobtools version 1.1.1 to remove potential contaminants (22). Blobtools requires three inputs– assembly, coverage, and hits. First, we mapped the raw reads back to the assembled genome using minimap2 version 2.20-r1061 with default settings to generate the coverage input (23). We then used the blastn module from NCBI BLAST+ version 2.12.0 to find sequence similarities between the assembled genome and 39 eukaryote and bacteria genomes (downloaded 25-October-2021; **Table S2**), which generated the hits input (24). Finally, we combined the assembly, coverage, and hits inputs using Blobtools version 1.1.1 to visualize and remove contaminant sequences. Contaminant sequences were classified as those sequences with abnormal coverage and GC proportions compared to the rest of the genome and having higher similarity with prokaryotic sequences.

### Omni-C scaffolding

InstaGRAAL was used for scaffolding the Dovetail Omni-C reads to the long-read contigs produced by the Flye assembler (25). The data was prepared with hicstuff v3.1.0, using BWA as the aligner, and the enzyme option was set to ‘mnase’ to be compatible with the OmniC data. Additionally, the filter setting was turned on to filter any short-range mapping events (26). Since no Omni-C-induced errors were detected, scaffolds were not improved by polishing.

### Characterization of repetitive sequences

Simple sequence repeats (SSRs) were identified with the R package micRocounter (27). The minimum number of repeated motifs to be considered an SSR was 6 for dinucleotides; 4 for trinucleotides; and 3 for tetra-, penta-, and hexanucleotides. A maximum gap to continue an SSR array was set to 1 nucleotide. Larger tandem repeats were identified using TRF v4.09.1 (28) with parameters 2, 5, 7, 80, 10, 50, and 2000. The maximum expected length of any repeat array was set to 10 Mbp. EDTA v2.0.0 (29) was used with the “sensitive” parameter set to 1 to construct a library and annotate interspersed repeats across the genome assembly. Identity with consensus sequence of transposable elements identified by homology was an output of EDTA.

R scripts incorporating ape (30) and SeqinR (31) were used to compile and split overlapping repeat annotations. Partially overlapping repeats were split 50/50. Fully overlapping repeat annotations were split 25/50/25, with the first and last 25% of the overlapping region attributed to the larger repeat. Plots were made using ggplot2 (32).

### Read coverage and synteny analysis

We calculated the normalized read coverage of each scaffold in RStudio and used the average genomic coverage to identify the scaffold that likely represents the X chromosome. To assess assembly quality, syntenic regions between the *D. frontalis* assembly and the new female assembly of *D. ponderosae* (13) were visualized with Circos v0.69-9 (33). Fasta files were cleaned using EDTA v2.0.0 (29). Scaffolds and contigs under 2Mb were removed using SeqKit (34) before creating the necessary karyotype files. Genome alignments were obtained using minimap2 v2.24 (23), and the resulting output file was then used to create a links file. This links file was used to generate the Circos plot. Conservation of Stevens elements were visualized using the same genome alignment data and the RIdeogram package (35).

### Transcriptome sequencing and assembly

cDNAs from SPB specimens were sequenced on an Illumina MiSeq instrument in using both 2×75bp and 2×150bp reads **(Table S3)**. Quality assessment of the data was performed using FastQC (36). TrimGalore (37) was used to remove reads from the dataset that are under the phred threshold of 30 and under the length threshold of 20bp. After low-quality reads were removed, contaminant sequences were identified using FastqScreen. The small size of the organism necessitated extracting RNA from whole-body samples, and contaminant sequences from the gut microbiome or SPB symbionts may have been present. After contaminants were identified, the RNAseq reads were mapped to contaminant genomes with the Burrows-Wheeler Alignment tool (38) and filtered according to map quality. rRNA contamination was also removed by mapping the RNAseq reads to a comprehensive set of Coleopteran rRNA sequences retrieved from the SILVA rRNA gene database (39). The remaining reads should represent only mRNA expressed by female, male, and larval SPB samples.

Transcriptome assembly was carried out using the Trinity *de novo* assembly pipeline (40). To remove redundancy, transcripts were subsequently clustered using the cd-hit-est tool available through the CD-HIT software package (41). The TransDecoder (42) pipeline, which leverage on BLAST (24) and Pfam (43) evidence, was used to identify transcripts in both the full and reduced assemblies that represent the longest open reading frame (ORF). TransDecoder filters out smaller isoforms and spurious or chimeric assemblies. The final draft assembly is a complete, non-redundant set of transcripts expressed by *D. frontalis*.

### Removal of transcripts containing transposable elements

To remove putative transcripts encoded by transposable elements (TEs) we performed a BLAST search of transcript sequences against the SPB library of TEs using the following modified parameters: -ungapped -max_hsps 5 -max_target_seqs 10 -evalue. The BLAST results were merged using the merge program in the bedtools suite (44), retrieving 1,893 transcripts with TEs content. The 1,070 transcripts with TE sequence coverage ≥50% were removed.

### Gene annotation

We used SPALN2 (45) to align the 40,493 transcripts onto the SPB genome assembly and mapped 39,588 transcripts, with the following parameters: -Q7 -O6 -t48 -d. To determine the number of loci we applied the program cluster in the bedtools suite (44) to exon and gene coordinates in the gff3 file, then identified for each cluster the main transcript by prioritizing ORF completeness (presence of both start and stop codons) and ORF length. Functional annotation of the 39,588 transcripts and the 13,354 loci was carried out using eggNOG-mapper v2 with default parameters (46).

### Gene family analysis

Genome assembly and protein fasta files and gff files of 17 gene sets and were obtained from the NCBI genome database **(Table S4**). The gff files were used to identify the longest coding sequence/protein per locus and the corresponding transcript IDs, in order to avoid including multiple isoforms/proteins in loci with alternative transcript data in gene family size analyses. Protein sequence files were filtered accordingly in order to include only the longest protein for each gene. These sequence files were used to infer gene families using OrthoFinder with default settings (47). Protein sequences of *Drosophila melanogaster* and *Dendroctonus ponderosae* (MPB) genes belonging to orthogroups of interest were used for functional enrichment analyses in STRING (48).

We inferred gene family expansions and contractions along the phylogeny of the 14 beetles and 3 outgroup species using CAFE 4 (49). Gene families with no variation across species, highly variable gene families (standard deviation>3) and families present in fewer than 6 species were removed, leaving a total of 8,903 orthogroups analyzed with CAFE. We ran the program with default parameters and one lambda, as we did not have specific hypotheses to test regarding variation in the rate of gene gain and loss along the species phylogeny. Sequence similarity searches to verify gene losses were performed using the standalone version ncbi-blast-2.11.0+ of BLAST+ (24) with default parameters except -max_hsps 10 -max_target_seqs 20 -ungapped - comp_based_stats F -evalue 0.1.

### Plant cell wall-degrading enzyme genes

Genes encoding for plant cell wall-degrading enzymes (PCWDEs) were identified in beetles by searching for the key words ‘Pectinesterase’ and ‘Glyco_hydro’ for carbohydrate esterases (CE) and glycoside hydrolases (GH), respectively, in the eggNOG-mapper annotation. Polysaccharide lyase (PL) genes were retrieved searching for ‘PL4’ in the CAZy database results in the eggNOG-mapper annotation. These genes were mapped onto the orthogroups from OrthoFinder. All genes from those orthogroups were then retrieved from the 17 analyzed species.

### Identification of transposable elements in gene sets

Protein domain names were retrieved using the PFMAs results from the eggNOG-mapper v2 analysis described above. We screened domain names using the following TE-associated domain keywords: DDE, hAT, integrase, RVT, MULE, Retrotrans, rve, gag, Tnp, Helitron, THAP. Protein sequences containing these domains were retrieved and used for local searches against the corresponding genomes using the ncbi-blast-2.11.0+ version of BLAST+. The BLAST parameters were set to default except for - max_hsps 10 -max_target_seqs 20 -ungapped -comp_based_stats F -evalue 0.1. As control, gene representatives from the twelve non-TE gene families with the highest average gene count across all species were also analyzed. Copy numbers for each gene were estimated by counting BLAST hits of at least 50 amino acids in non-overlapping genomic regions with the four possible combinations of distance between hits 20kb or 50kb and percentage identity 50% or 75%.

## Results and Discussion

### Genome assembly

The final scaffolded assembly contained 381 scaffolds at a total length of 173.7 Mb, with a scaffold N50 of 24.8 MB and 97.72% of the assembled genome localized in nine scaffolds larger than 50 kb. BUSCO analyses of the scaffolded assembly contained 94.2% of the 2,124 Endopterygota conserved orthologs as complete copies and 1.2% of genes with additional copies.

### Repetitive sequence identification

Almost 30% of the SPB assembled genome was identified as repetitive. Chromosome-level scaffolds contained a lower proportion of repeats compared to small scaffolds, as expected given the challenges posed by repetitive DNA to the assembly of long pseudomolecules (**Fig. 1**).

**Figure 1.**
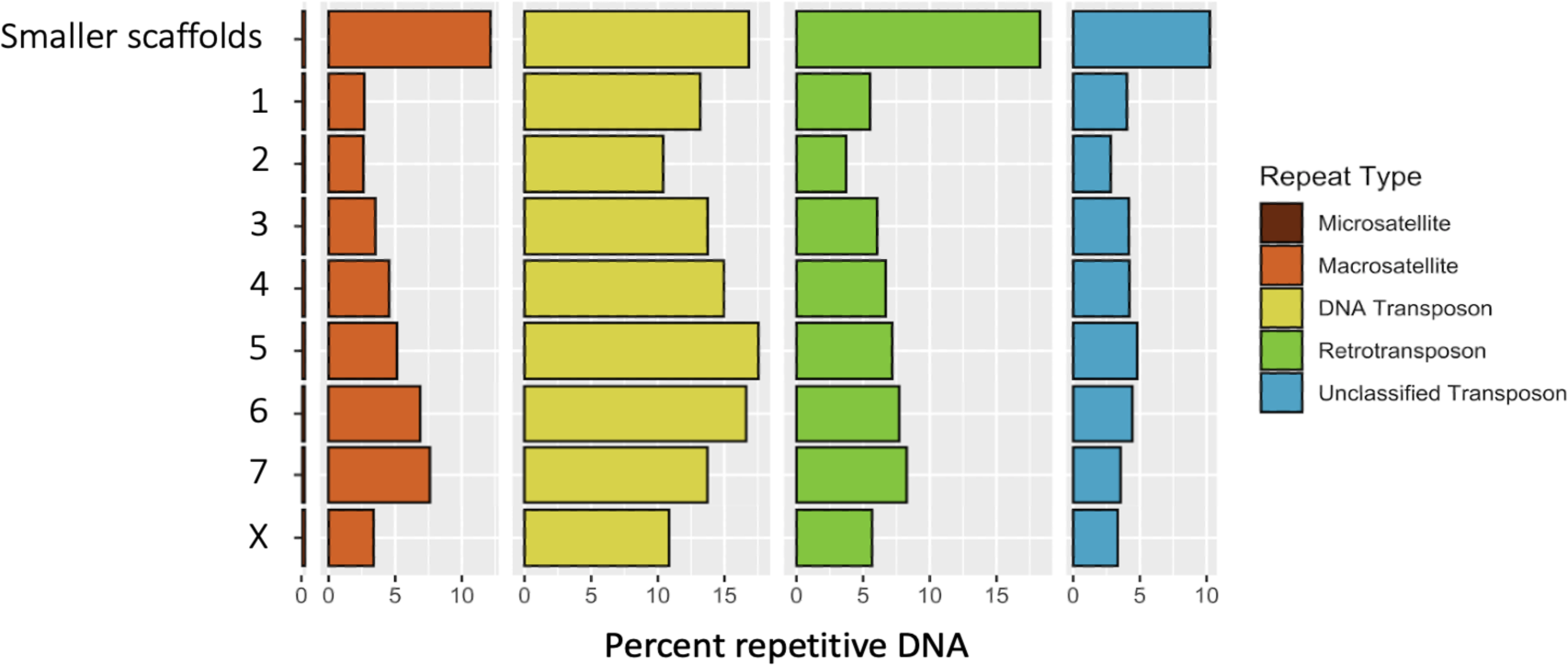
Distribution of repetitive elements. This figure displays the distribution of repetitive elements in the genome assembly. The x-axis represents scaffold numbers, while the y-axis displays the percentage of each repetitive element type.

Transposable elements formed ∼23% of the assembled genome, similarly to what found in MPB (12, 50) but lower than in the larger genome of *D. valens* (14). Approximately 13% and ∼6% of the SPB genome was formed by DNA transposons and retrotransposons, respectively (**Table S5**). We also identified 71,000 tandem repeat arrays contributing ∼5% of the assembled genome. The telomeric (TTAGG)n repeat found in some members of Scotylinae (51) was not found at the termini of large scaffolds of the assembled genome.

### Synteny conservation with MPB and identification of the putative chromosome X in SPB

After normalizing read coverage across the genome assembly of *D. frontalis* for female and male samples, we found a reduction in male read coverage in scaffold 8, suggesting that this scaffold represents the X chromosome (**Fig. 2**).

**Figure 2.**
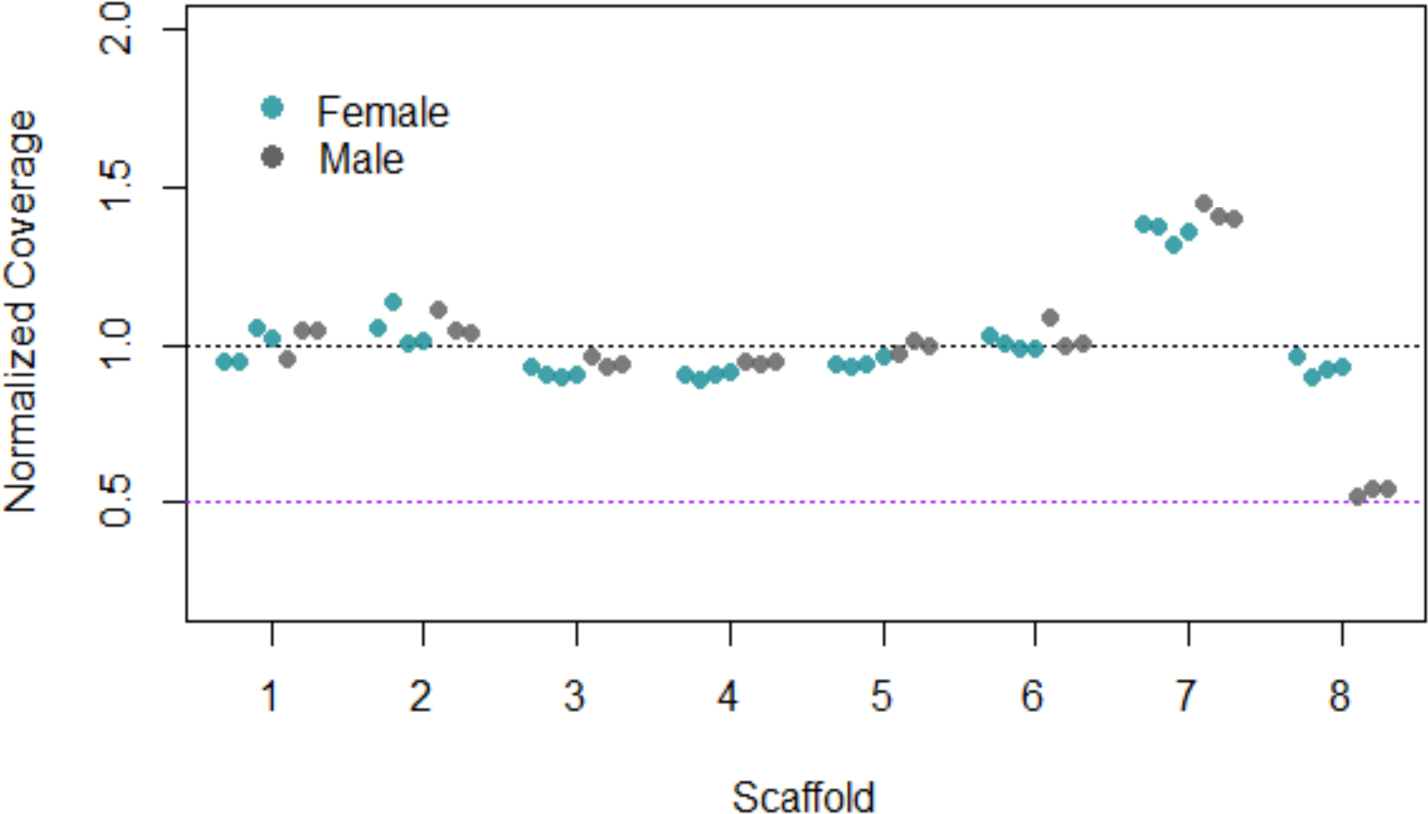
Normalized read coverage across the *D. frontalis* genome assembly. The teal points represent the normalized read coverage for female samples, while the gray points represent the corresponding coverage for male samples.

The comparison between the SPB and *D. ponderosae* genomes revealed high level of synteny conservation. The scaffold containing the X chromosome in SPB maps to scaffold 1 in *D. ponderosae*, which correspond to the neoXY system in *D. ponderosae* (**Fig. 3**). form a set of genomic regions with conserved synteny across multiple beetle species.

**Figure 3.**
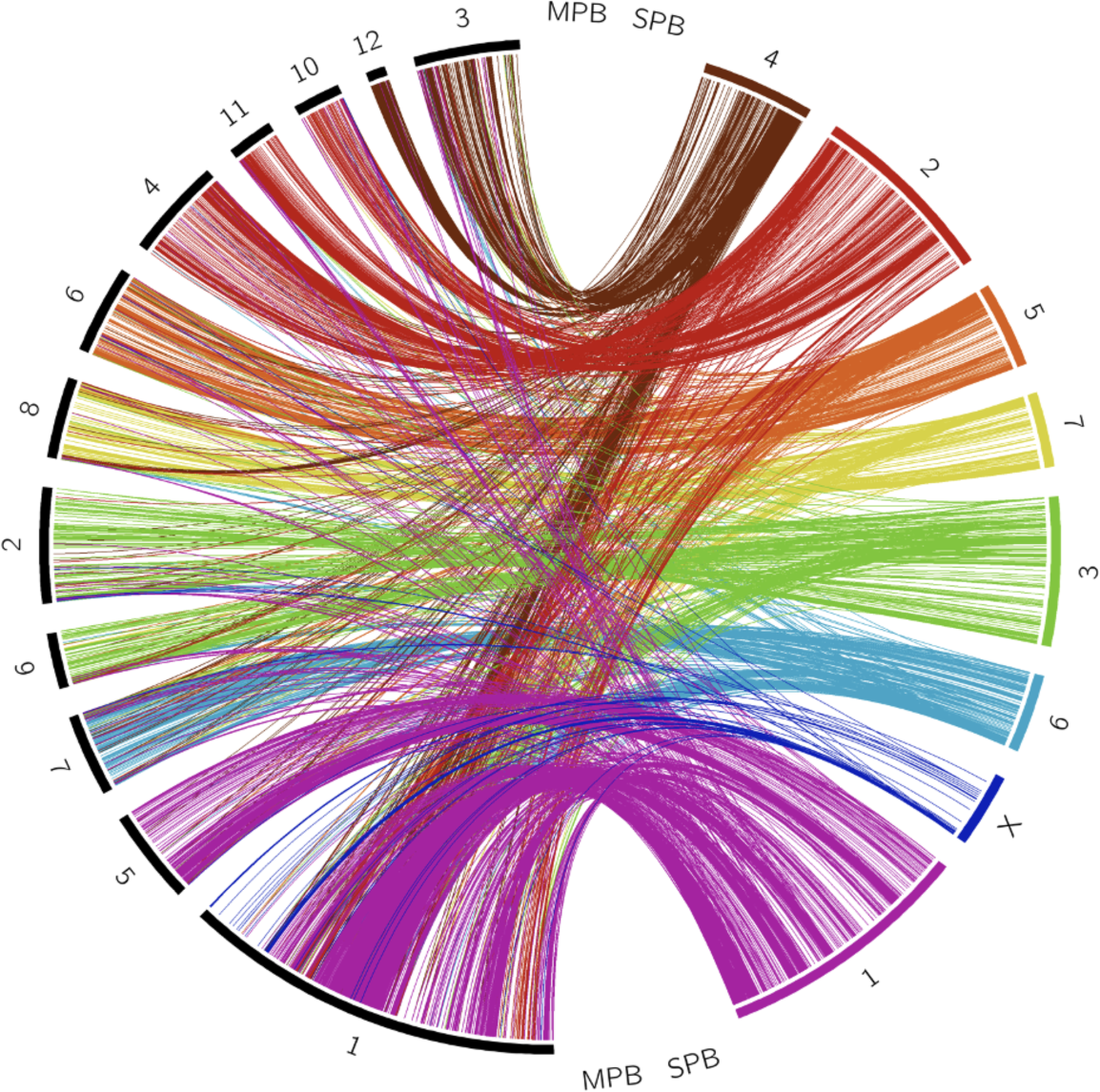
Synteny correspondence between *Dendroctonus frontalis* and *Dendroctonus ponderosae* chromosomes. The putative *D. frontalis* X chromosome shows synteny conservation with part of the neoX chromosome in *D. ponderosae*. *D. frontalis* chromosomes: colored segments. *D. ponderosae* chromosomes: black segments.

Based on the synteny analysis of *Tribolium castaneum* and five other beetles, Bracewell and colleagues have recently proposed that the Coleopteran ancestral karyotype was formed by nine Stevens elements, named after Nettie Stevens (52). A synteny plot based on SPB, *D. ponderosae* and *T. castaneum* genomes shows conservation of the nine Stevens elements (**Fig. 4**).

**Figure 4.**
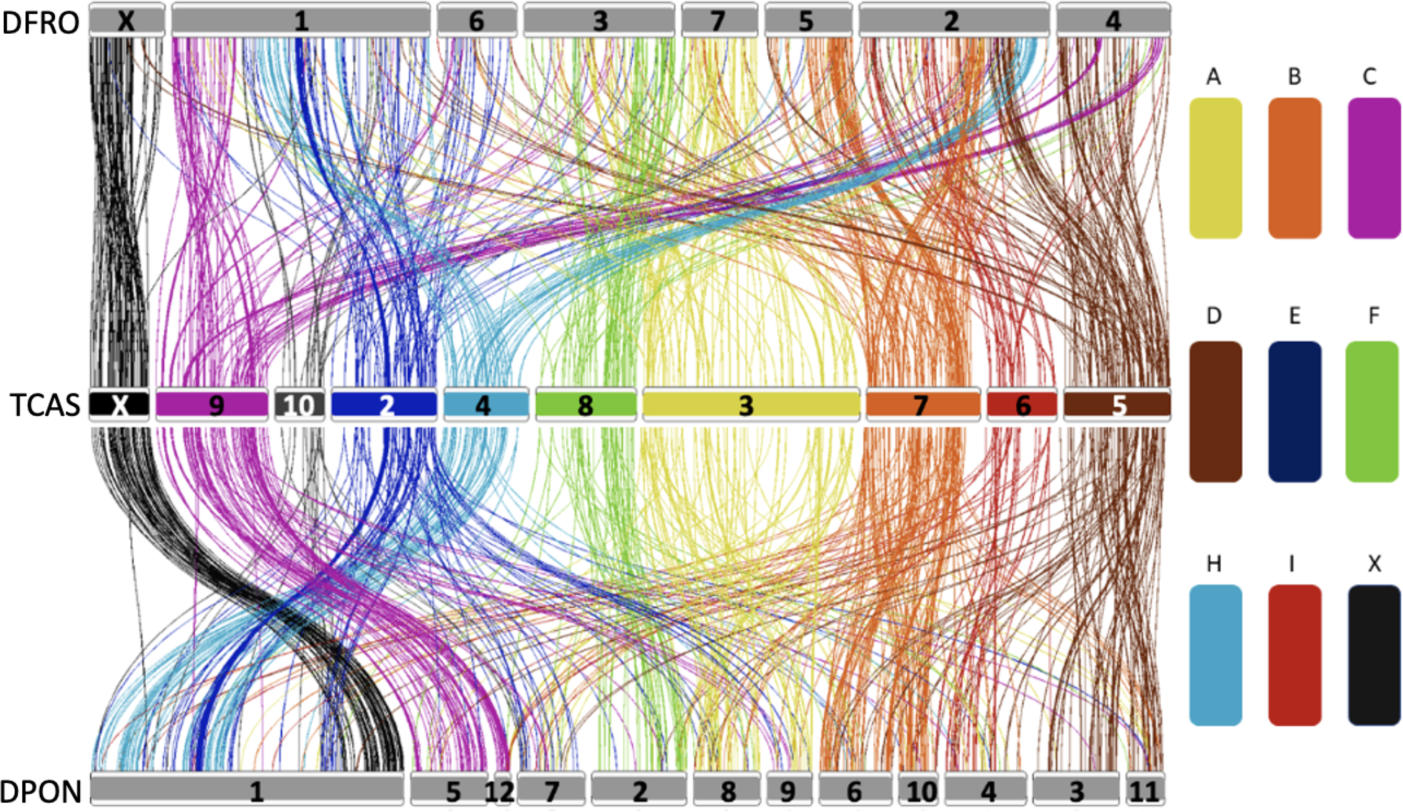
Synteny plot displaying conservation of Stevens elements. The conservation of the nine Stevens elements in three beetle genomes (top: *D. frontalis*, middle: *T. castaneum*, bottom: *D. ponderosae*).

### Gene annotation

We used a custom gene annotation pipeline to assemble transcripts from RNA-seq data and map them onto the genome assembly (see **Materials & Methods**). We identified 39,588 transcripts from 13,354 putative gene loci in the SPB genome (**Table 1**). A similar number of loci was identified in the recently improved assembly of *D. ponderosae* (Keeling et al. 2022) and in the *D. valens* genome (14), whereas the genome of *H. hampei* and *I. typographus* contains a significantly higher number of genes (15–17). Among all mapped transcripts, 9,678 (72.5%) were functionally annotated with eggNOG-mapper v2 (46) **(Table S6)**.

**Table 1.**
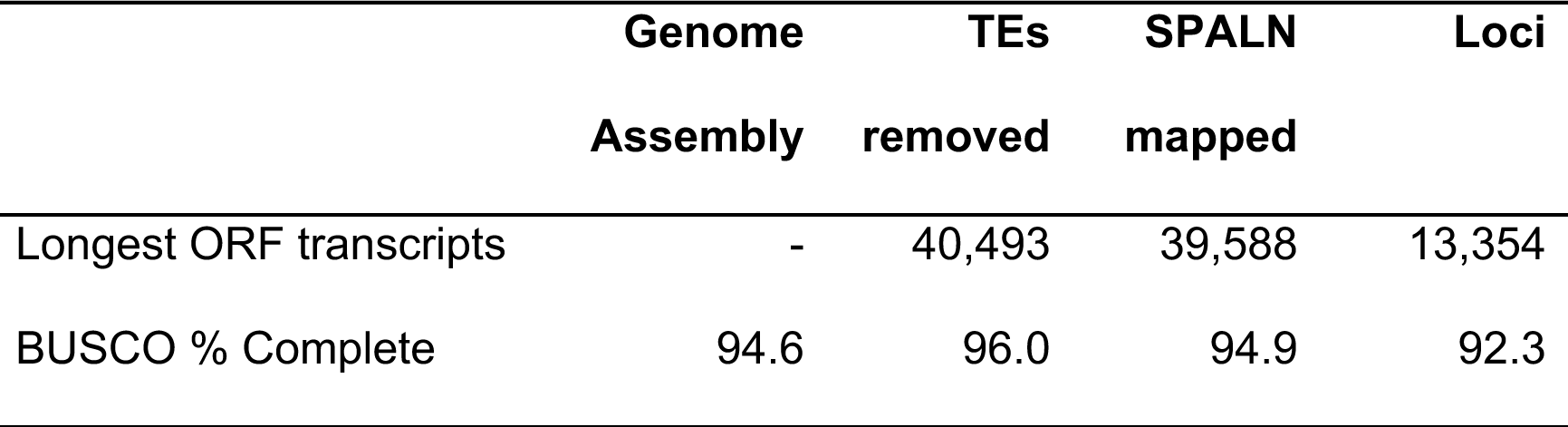
SPB genome and transcriptome metrics and gene annotation.

### Gene family analyses

We investigated changes in the gene content that might be associated with trait evolution in SPB and *Dendroctonus*. Leveraging on high quality genomic resources available for two non-SPB *Dendroctonus* species, two additional Scolytinae, nine other beetles and three non-beetle insects (**Table S4**), we built and analyzed orthogroups (gene families) using OrthoFinder (47). We identified 17,135 gene families present in at least two species and a high percentage of genes grouped in families in beetles (85-100%) and outgroup species (71-85%) **(Table S4)**.

To comprehensively examine gene family evolution in SPB and other *Dendroctonus* species, we analyzed gains and losses in orthogroups along the phylogeny of the fourteen beetles and three outgroup species using CAFE (49).

In SPB, *Dendroctonus* and Scolytinae, gene family contractions generally outnumbered expansions, with the exceptions of *Ips* and of the ancestral *Dendroctonus* branch **(Fig. 5)**. This is in agreement with the observed diminished gene content in *Dendroctonus* and increased gene count in *Ips* compared to other beetles. We next explored changes in gene family size in SPB, the ancestral branch of SPB and MPB, and the ancestral *Dendroctonus* lineage that could be relevant to the *D. frontalis* trait evolution and to the prevalence of the tree killing habit species in this genus of bark beetles.

**Figure 5.**
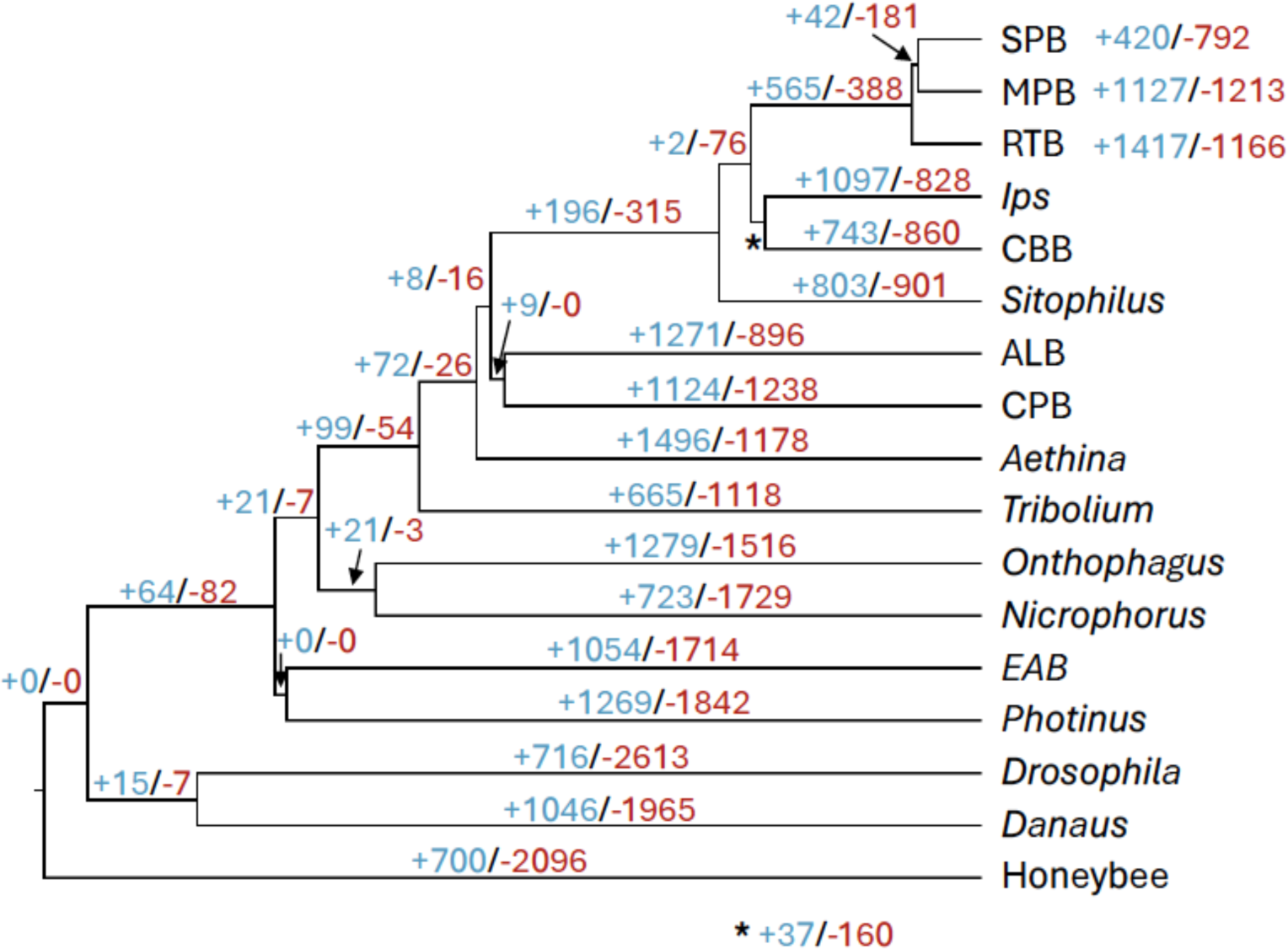
Gene family (orthogroups) changes along the phylogeny of fourteen beetles and three outgroup insects inferred using CAFE. The number of gene families with gains and losses are shown in blue and red, respectively. SPB: southern pine beetle (*Dendroctonus frontalis*). MPB: mountain pine beetle (*D. ponderosae*). RTB: red turpentine beetle (*D. valens*). *Ips*: *Ips typographus*. CBB: coffee berry borer (*Hypothenemus hampei*). *Sitophilus*: *Sitophilus oryzae*. ALB: Asian long-horned beetle (*Anoplophora glabripennis*). CPB: Colorado potato beetle (*A*). *Aethina*: *Aethina tumida*. *Tribolium*: *Tribolium castaneum*. *Onthophagus*: *Onthophagus taurus*. *Nicrophorus*: *Nicrophorus vespilloides*. EAB: emerald ash borer (*Agrilus planipennis*). *Photinus*: *Photinus pyralis*. *Drosophila*: *Drosophila melanogaster*. *Danaus*: *Danaus plexippus*. Honeybee: *Apis mellifera*.

**Figure 6.**
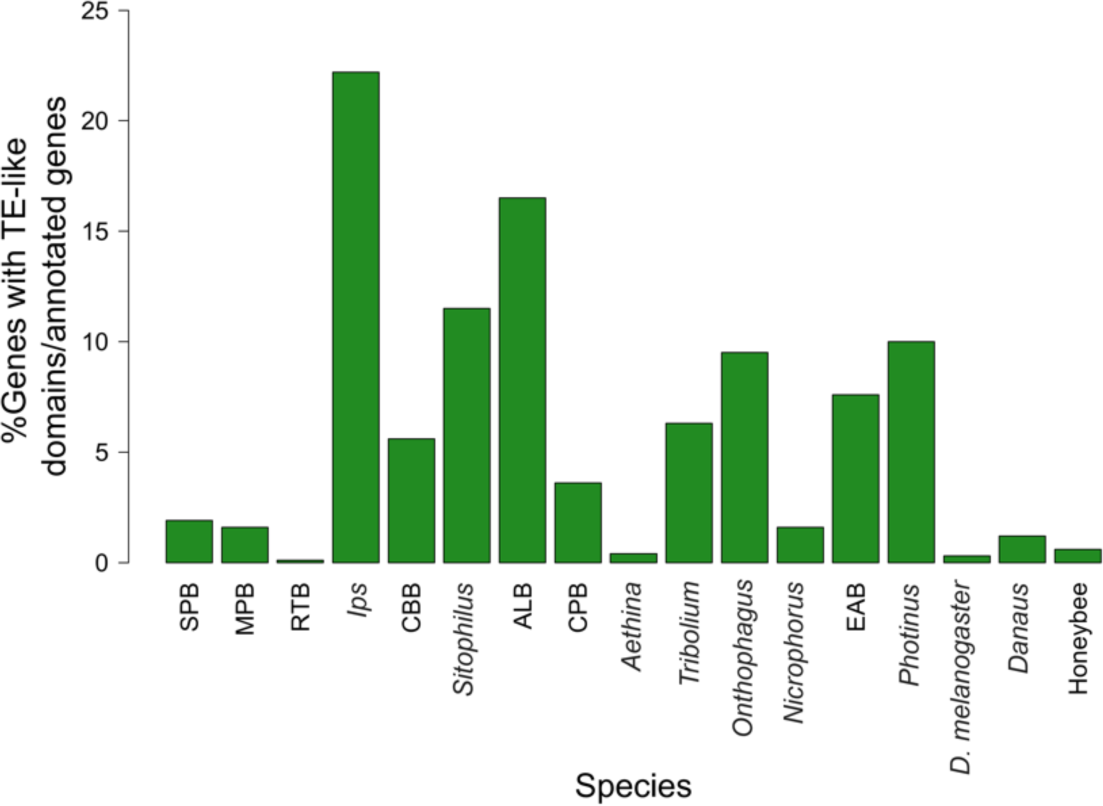
Proportion of functionally annotated genes encoding proteins with TE-derived domains. Name abbreviations are listed in **Table S4**.

The CAFE analysis revealed 792 families contracted in SPB; of these, 437 were apparently completely lost. Gene losses contribute significantly to species evolution and adaptation (53, 54), but remain poorly investigated in insects. To better assess the magnitude and potential biological impact of gene loss in SPB, we carried out BLAST searches of MPB genes belonging to these families against the SPB genome assembly and confirmed the lack of any homology for eight orthogroups. As BLAST analyses were permissive in order to retrieve SPB regions with low homology to the MPB genes, these results could include false positives (see also **Materials & Methods**).

Nevertheless, these indicate that several genes might still be unreported or only partially annotated in the SPB gene set; alternatively, they might have incurred pseudogenization in SPB. Genes missing in SPB included the serine/threonine-protein kinase *Tricornered*, a splicing factor, an amino acid transporter and a protein tyrosine phosphatase **(Table S7).**

Among the 420 orthogroups with gene gains in SPB, we examined the possible function of 293 families with more genes in SPB than other Scolytinae. Functional enrichment was determined using MPB and *D. melanogaster* members of these families (one gene per family) using the STRING database (48). Genes encoding for membrane proteins and extracellular matrix proteins experience high rates of duplication in SPB, suggesting a key role of proteins at the cellular-environment interface in adaptation and specialization **(Table S8)**. Results using MPB genes produced only one significant enrichment, which is expected given the limited functional annotation of most MPB genes, corresponding to the large ‘Cell periphery’ cellular component **(Table S8)**. To further dissect the contribution of gene duplication to trait evolution in SPB, we analyzed a subset of 85 orthogroups that contained two genes in SPB and one gene in other Scolytinae and in *D. melanogaster* (subset ‘2-to-1 SPB’). Three large partially overlapping networks, ‘Plasma membrane bounded cell projection organization’, ‘Cell morphogenesis involved in differentiation’ and ‘Animal organ development’, stood out as processes with significant gene family expansions in SPB **(Table S8)**.

We next investigated contractions and expansions of gene families in the ancestral branch of SPB and MPB. A total of 141 gene families appeared extinct in both SPB and MPB compared to their sister *Dendroctonus* species RPB. BLAST searches revealed homologous hits in the SPB and MPB genomes for all but 11 of these 141 orthogroups. Several genes conserved across beetles and other insects were lost in SPB and MBP, including a locus required for the development of *D. melanogaster* ovarian follicles (*Kuduk*) and a gene regulating tube morphogenesis in the tracheal system (*Ccm3*) **(Table S7)**. Notably, *Ccm3* genetically interact with *Tricornered* (55), one the genes uniquely lost in SPB, implying significant changes in the control of tube morphogenesis in the *Dendroctonus* clade. Lineage-specific gene duplications in the SPB/MPB clade occurred only in 42 gene families but showed a significant enrichment for ‘Cell junction’ and ‘Mitotic spindle’ processes **(Table S8)**.

Along the *Dendroctonus* stem lineage, we retrieved 388 orthogroups with contractions, including 218 completely lost gene family. BLAST analyses using *Ips*, CBB and *D. melanogaster* genes belonging to these families confirmed the loss of 17 orthogroups. The 23 *D. melanogaster* genes with no homologs in *Dendroctonus* contained highly conserved loci involved in survival to dietary restriction and oxidative stress (*Thor*), maintenance of the female germ line (*Stonewall* and *Brickwall*), transcription of mitochondrial proteins (*Spargel*), mitotic chromosome condensation (*prod* and *Mink*), and repair of UV-induced DNA damage (*phr*) **(Table S7)**.

A total of 565 gene families showed gene gains in the *Dendroctonus* clade. We searched for functional enrichments in the 200 expanded families with the largest increase between the *Dendroctonus* and Scolytinae branches. Gene family expansions were associated to a variety of processes that might be involved in adaptation, including ‘Response to stimulus’, ‘Locomotion’ and ‘Compound eye development’ **(Table S8)**.

### Plant cell wall-degrading enzymes genes in SPB

Plant cell wall-degrading enzymes (PCWDEs) are required to digest cellulose, pectin and other complex carbohydrates that constitute the plant cell wall, a major energy source for herbivorous insects (56). A recent survey of Coleopteran transcriptomes and genomes showed correlated expansions in horizontally acquired PCWDEs with adaptive radiations and specialized herbivory (57). PCWDE family expansions were particularly common among Phytophaga and Buprestoidea, the most taxonomically diverse and specialized lineages within Coleoptera. Using Pfam domain results we predicted a total of 651 PCWDE genes across the 17 species (**Table S9**). We found very similar number of PCWDE genes in the wood-boring species MPB, ALB and EAB compared to those previously described by McKenna et al. (2019). Our novel annotation of PCWDE genes in SPB and RTB confirmed a high number of these genes across *Dendroctonus*, albeit to a less extent than observed in MPB (**Table S9**).

### Gene content reduction in *Dendroctonus* and gene misannotation in beetles

The three sequenced *Dendroctonus* species contain less than 14,000 genes, fewer than all the other beetles included in these analyses **(Table S4)**. Notably, 43-79% more genes have been reported in the two other sequenced Scolytinae genomes, the spruce bark beetle and the coffee berry borer, than in *Dendroctonus*. We sought to disentangle the possible contribution of biological factors and gene annotation shortfalls to the diminished gene repertoire in *Dendroctonus* genomes.

First, we found a lower gene annotation completeness between *Dendroctonus* and other species in the suite of highly conserved Endopterygota genes assessed by BUSCO, confirming the gene family analysis results indicating loss of several conserved genes in *Dendroctonus* genomes **(Table S10)**. *Dendroctonus* showed on average ∼1% of more missing conserved genes than other beetles, or about 160 loss genes after extrapolating to an overall beetle gene count of ∼17,000.

However, the BUSCO analysis is limited to a subset of genes that are unlikely to be representative of the overall gene complement of a species, particularly for gene families with high rates of turnover. Therefore, we expanded our analyses of gene family size to better determine how gene gains and losses shaped the gene content differences across beetles. The CAFE results showed that gene gain and loss rates were similar in *Dendroctonus* compared to other species **(Fig. S1)**. Moreover, the average size of orthogroups used in the CAFE analysis is nearly identical between *Dendroctonus* and other species **(Table S11)**. Thus, we reasoned that gene content differences among these two groups of beetles must lie within the 9,865 gene families present in beetles that were excluded from the CAFE analyses. Among these, the 3,764 families occurring in *Dendroctonus* showed no difference in size between the two groups of beetles **(Table S11)**. This suggests that gene content is higher in non-*Dendroctonus* species primarily due to orthogroups that do not occur in the *Dendroctonus* clade. Notably, nearly 83% of these orthogroups are present in less than three beetle genomes, indicating that they derive from the emergence of novel lineage-specific genes (**Tables S4**). Additionally, only ∼228 genes in *Dendroctonus* were not included in orthogroups, compared to ∼1,256 genes in non-*Dendroctonus* species, supporting the higher proportion of lineage-specific genes in the latter group **(Table S11).**

We next sought to assess if the high number of lineage-specific genes in non-*Dendroctonus* beetles could be caused by assembly and annotation artifacts (58). In particular, we investigated the potential role of transposable elements (TEs) as a source of gene annotation artifacts. Insect genomes harbor several genes that originated from the ‘domestication’ of TEs (59), but they typically form a small portion of the overall gene repertoire and should not account for major differences in gene counts between species. We developed a novel approach to rapidly screen the gene sets of all analyzed species for the presence of an excess of TEs-derived genes. First, we identified proteins containing domains derived from TEs based on eggNOG-mapper annotation and searchers of TE-associated keywords. We observed a much higher numbers of genes containing TE-derived domains in the bark beetle species *Ips* and CBB compared to *Dendroctonus*, as well as in ALB, *Sitophilus*, *Onthophagus*, and *Photinus* **(Fig. 3; Table S12)**.

To verify if most of these genes represent misannotated TEs, we performed BLAST searches for each candidate TE-derived protein against their genome of origin and estimated their copy number using several combinations of sequence identity and distance between genomic hits (**see Materials & Methods**). The same approach was used to estimate the copy number of the 12 largest gene families in our dataset that do not contain TE-derived domains. Putative genes with TE domains had more copies on average than non-TE genes in every species, with *Ips* having the highest copy number for the former **(Table S12)**. These estimates might have been slightly inflated for non-TE genes due to multiple hits for the same gene within the distance range between hits. Additionally, we found a higher proportion of copies with stop codons between genes with TE domains compared to other genes in each species for most comparisons **(Table S12)**. This is expected for misannotated TEs, as many copies of transposable elements contain disabled coding sequences.

Altogether, these results suggest that the diminished gene content in the genus *Dendroctonus* is due to a combination of high levels of gene loss in *Dendroctonus* species and a large expansion of lineage-specific gene family repertoires in many non-*Dendroctonus* beetle, at least partly due to TE misannotation.

### Conclusion

Genome sequencing and analysis efforts are essential to identifying the genetic basis of pest behavior in insects and to inform advanced pest management strategies. Using long-read genome sequencing and high-throughput transcriptomic data, we generated a chromosome-level assembly and high-quality gene annotation of the southern pine beetle *Dendroctonus frontalis*, a major conifer pest. We confirmed the extensive synteny conservation across beetles and identified the putative X chromosome in SPB. Gene family analyses of 14 beetle species revealed several losses of conserved genes and lineage-specific gene gains in SPB and other *Dendroctonus* species. Overall, the *Dendroctonus* clade experienced numerous gene losses and a reduced rate of formation of novel gene families, which seem to account for the diminished gene complement in this genus. However, we found strong evidence of widespread misannotation of transposable elements in the gene complement of many non-*Dendroctonus* beetles, which appear to contribute significantly to the observed variation in gene content in Coleoptera.

## Data Availability

The *Dendroctonus frontalis* genome assembly sequence is available through the NCBI BioSample ID PRJNA1100959. Raw transcriptome sequencing reads are available through the SRA ID PRJNA1102401. Datasets and gene family analysis results are available through the following Figshare repository: https://doi.org/10.6084/m9.figshare.25491793.v1.

## Acknowledgments

We thank the Eppley Foundation for Research, the Texas A&M AgriLife Research and the Texas A&M Forest Service for supporting this project. CC was supported by the USDA National Institute of Food and Agriculture project 1019860; HB was supported by the National Institute of General Medical Sciences at the National Institutes of Health R35GM138098. We are grateful to Brian T. Sullivan for assistance with collecting *D. frontalis* specimens.

