## Supplementary material for "Genome assembly of the southern pine beetle (*Dendroctonus frontalis* Zimmerman) reveals the origins of gene content reduction in *Dendroctonus*": Fig. S1

**Title: Fig. S1**

**Legend: Figure S1**. Gene gains (blue) and losses (red) along the phylogeny of fourteen beetles and three outgroup insects inferred using CAFE. SPB: southern pine beetle (*Dendroctonus frontalis*). MPB: mountain pine beetle (*D. ponderosae*). RTB: red turpentine beetle (*D. valens*). *Ips*: *Ips typographus*. CBB: coffee berry borer (*Hypothenemus hampei*). *Sitophilus*: *Sitophilus oryzae*. ALB: Asian long-horned beetle (*Anoplophora glabripennis*). CPB: Colorado potato beetle (*A*). *Aethina*: *Aethina tumida*. *Tribolium*: *Tribolium castaneum*. *Onthophagus*: *Onthophagus taurus*. *Nicrophorus*: *Nicrophorus vespilloides*. EAB: emerald ash borer (*Agrilus planipennis*). *Photinus*: *Photinus pyralis*. *Drosophila*: *Drosophila melanogaster*. *Danaus*: *Danaus* *plexippus*. Honeybee: *Apis mellifera*.

**Title: Table S1**

**Legend: Table S1. Collected SPB speciments.**

**Title: Table S2**

**Legend: Table S2. Species used to detect possible contamination in the SPB genome assembly**

**Title: Table S3**

**Legend: Table S3. RNA-seq data.**

**Title: Table S4**

**Legend: Table S4. Genome, gene and gene family features of 14 beetles and 3 insect outgroups used in gene family analyses.**

**Title: Table S5**

**Legend: Table S5. Transposable element content in the SPB genome.**

**Title: Table S6**

**Legend: Table S6. eggNOG-mapper annotation of SPB genes.**

**Title: Table S7**

**Legend: Table S7. Gene losses in the *Dendroctonus* clade.**

**Title: Table S8**

**Legend: Table S8. Functional enrichment results from STRING analyses.**

**Title: Table S9**

**Legend: Table S9. Annotation of PCWDE genes in 14 beetles and 3 insect outgroup species.**

**Title: Table S10**

**Legend: Table S10. BUSCO results of 17 species.**

**Title: Table S11**

**Legend: Table S11. Gene family comparisons between *Dendroctonus* and other beetles.**

**Title: Table S12**

**Legend: Table S12. Genes with TE-associated domains, estimated copy numbers and stop codons/copy. TE-domain results are shared in blue, gene results are shared in gray.**
